## Supplementary_notes for "Power analysis of transcriptome-wide association study: implications for practical protocol choice"

### Supporting Information

#### Abbreviations & Notation table

GWAS: Genome-wide association study;

TWAS: Transcriptome-wide association study;

MAF: Minor allele frequency;

$X$: Genotype

$Z$: Expression

$Y$: Phenotype

$v$: NCP

$\beta$: coefficient for a fixed effect term

$u$: random effect term in a mixed model

$h^2$: heritability or variance component coefficient

$\sigma^2$: variance component to be estimated in a mixed model

#### Availability of data and materials.

The 1000 Genomes Project genotype data is downloaded from the project website:

https://www.internationalgenome.org

All the code to conduct the analysis is in our GitHub:

https://github.com/boweiding/power-analysis-of-TWAS

#### S1 Appendix: Solving the ElasticNet model for predicting expression using genotype

In our study, we optimize tuning parameters through a 10-fold cross-validation. First we set up a sequence of candidate $\alpha$ values with $\alpha \in (0.1, 0.15, 0.2, 0.25, \cdots, 0.9)$, under each $\alpha$, we conduct the 10-fold cross-validation to select the optimal $\lambda$ value which minimizes the objective function. For 10-fold cross-validation, the sample is randomly divided into 10 folds of equal size. Then the model is fit 10 times under a specific $\lambda$ with each time omitting one of the folds without repetition for estimating parameters. For each time, the objective function will be computed in the omitted fold (test set). Then the mean objective function is the mean of the values among ten different test sets. Therefore, the optimal $\lambda$ is the one under which the objective function is minimized among the whole sequence of $\lambda$ under a specific $\alpha$. After obtaining an optimal $\lambda$ under each $\alpha$, we compare the objective function of each combination of $\lambda$ and $\alpha$ and choose the minimized error among all combinations. Thus, the tuning parameters $\lambda$ and $\alpha$ with minimized objective function are the optimal ones.

#### S2 Appendix: Solving the variance component $\sigma_g^2$

In the decorrelation matrix $D_x$ given by (10), although the kinship matrix $K_x$ is known and further $\Lambda_x$ and $U_x$ can be decomposed from $K_x$, the variance components $\sigma_g^2$ and $\sigma_e^2$ are unknown parameters and should be estimated in order to obtain an estimated decorrelation matrix. The estimation of variance components $\sigma_g^2$ and $\sigma_e^2$ is generally mature and accurate in practice when sample size is at the level of thousands and the effect of each SNP or expression on the phenotype is small [35,42].

To estimate $\sigma_g^2$ and $\sigma_e^2$, following the efficient mixed-model association eXpedited (EMMAX) [35], we use a variance component model

$$Y_c = u + \varepsilon \tag{19}$$

to obtain the maximum likelihood estimator (MLE) of $\sigma_g^2$ and $\sigma_e^2$, where $u$ and $\varepsilon$ are given in (1.2), $Y_c = Y - \bar{Y}\mathbf{1}$ is an $n \times 1$ vector, $Y$ is the $n \times 1$ phenotype vector, and $\bar{Y}$ is the sample mean $\bar{Y} = \sum_{i=1}^{n} Y_i$. Here $Y_c$ is essentially the centralized $Y$ with sample mean zero in order to have model (19) being appropriate [35]. In order to facilitate our following derivation and calculation, we define

$$\delta = \frac{\sigma_e^2}{\sigma_g^2}. \tag{20}$$

Now it holds approximately that $Y_c \sim N\left(0, \sigma_g^2(K_x + \delta I)\right)$. The next claim presents the MLEs of $\sigma_g^2$ and $\sigma_e^2$.

**Claim 1.** *Under models (19) and (20), the MLEs of $\sigma_g^2$ and $\sigma_e^2$ are given by*

$$\hat{\sigma}_g^2 = \frac{1}{n}\sum_{i=1}^{n} \frac{V_{xi}^2}{\lambda_{xi} + \hat{\delta}}, \tag{21}$$

$$\hat{\sigma}_e^2 = \frac{\hat{\delta}}{n}\sum_{i=1}^{n} \frac{V_{xi}^2}{\lambda_{xi} + \hat{\delta}}, \tag{22}$$

*where $V_x = U_x^T Y_c = (V_{x1}, \cdots, V_{xn})^T$ and $\hat{\delta}$ is the solution to the estimating equation*

$$\sum_{i=1}^{n}\left[\frac{nV_{xi}^2}{\left(\sum_{i=1}^{n} \frac{V_{xi}^2}{\lambda_{xi} + \delta}\right)\left(\lambda_{xi} + \hat{\delta}\right)^2} - \frac{1}{\lambda_{xi} + \hat{\delta}}\right] = 0. \tag{23}$$

*Proof*. Let $\Sigma = \sigma_g^2(K_x + \delta I)$, then by the eigen-decomposition $K_x = U_x \Lambda_x U_x^T$ and the facts that $U_x U_x^T = U_x^T U_x = I$ we have

$$\begin{aligned} U_x^T \Sigma U_x &= \sigma_g^2 U_x^T K_x U_x + \sigma_g^2 \delta U_x^T U_x \\ &= \sigma_g^2(\Lambda_x + \delta I) \\ &= \begin{pmatrix} \sigma_g^2(\lambda_{x1} + \delta) & 0 & 0 & \cdots & 0 \\ 0 & \sigma_g^2(\lambda_{x2} + \delta) & 0 & \cdots\cdots & 0 \\ \cdots & \cdots & \cdots & \cdots\cdots & \cdots \\ 0 & 0 & 0 & \cdots\cdots & \sigma_g^2(\lambda_{xn} + \delta) \end{pmatrix}, \end{aligned}$$

and then

$$|\Sigma| = |U_x^T \Sigma U_x| = \prod_{i=1}^{n} \sigma_g^2 \, (\lambda_{xi} + \delta).$$

With $V_x = U_x^T Y_c$, we rewrite $Y_c^T \Sigma^{-1} Y_c$ as

$$\begin{aligned} Y_c^T \Sigma^{-1} Y_c &= (U_x^T Y_c)^T U_x^T \Sigma^{-1} U_x U_x^T Y_c \\ &= V_x^T (U_x^T \Sigma U_x)^{-1} V_x \\ &= \sum_{i=1}^{n} V_{xi}^2 \left[\sigma_g^2(\lambda_{xi} + \delta)\right]^{-1}. \end{aligned}$$

Now the log-likelihood function of $\sigma_g^2$ and $\delta$ is given by

$$
\begin{aligned}
l(\sigma_g^2, \delta) &= \ln[f(Y_{c1}, Y_{c2}, \cdots, Y_{cn})] \\
&= \ln\left[(2\pi)^{-\frac{n}{2}}|\Sigma|^{-\frac{1}{2}} \exp\left(-\frac{1}{2} Y_c^T \Sigma^{-1} Y_c\right)\right] \\
&= -\frac{n}{2}\ln(2\pi) - \frac{1}{2}\ln|\Sigma| - \frac{1}{2} Y_c^T \Sigma^{-1} Y_c \\
&= -\frac{n}{2}\ln(2\pi) - \frac{1}{2}\ln\left[\prod_{i=1}^{n} \sigma_g^2\,(\lambda_{xi} + \delta)\right] - \frac{1}{2}\sum_{i=1}^{n} V_{xi}^2\,[\sigma_g^2(\lambda_{xi} + \delta)]^{-1} \\
&= -\frac{n}{2}\ln(2\pi) - \frac{n}{2}\ln\sigma_g^2 - \frac{1}{2}\sum_{i=1}^{n}\ln(\lambda_{xi} + \delta) - \frac{1}{2}\left(\sigma_g^2\right)^{-1}\sum_{i=1}^{n} V_{xi}^2\,(\lambda_{xi} + \delta)^{-1}.
\end{aligned}
$$

Then the MLE estimating equations for $\sigma_g^2$ and $\delta$ are

$$
\frac{\partial}{\partial(\sigma_g^2)} l(\sigma_g^2, \delta) = -\frac{n}{2\sigma_g^2} + \frac{1}{2\sigma_g^4}\sum_{i=1}^{n}\frac{V_{xi}^2}{\lambda_{xi} + \delta} = 0, \tag{24}
$$

$$
\frac{\partial}{\partial\delta} l(\sigma_g^2, \delta) = -\frac{1}{2}\sum_{i=1}^{n}\frac{1}{\lambda_{xi} + \delta} + \frac{1}{2}\sum_{i=1}^{n}\frac{V_{xi}^2}{\sigma_g^2(\lambda_{xi} + \delta)^2} = \frac{1}{2}\sum_{i=1}^{n}\left[\frac{V_{xi}^2}{\sigma_g^2(\lambda_{xi} + \delta)^2} - \frac{1}{\lambda_{xi} + \delta}\right] = 0. \tag{25}
$$

From (24) we obtain

$$
\sigma_g^2 = \frac{1}{n}\sum_{i=1}^{n}\frac{V_{xi}^2}{\lambda_{xi} + \delta},
$$

and plugging this into (25) we obtain

$$
\sum_{i=1}^{n}\left[\frac{nV_{xi}^2}{\left(\sum_{i=1}^{n}\frac{V_{xi}^2}{\lambda_{xi} + \delta}\right)(\lambda_{xi} + \delta)^2} - \frac{1}{(\lambda_{xi} + \delta)}\right] = 0.
$$

Finally, by (20), hence the result.

It has been shown [35] that the MLEs of $\sigma_g^2$ and $\sigma_e^2$ given in (2.4) and (2.5) respectively have good accuracy. In order to solve for $\delta$ in (2.6), one may use Newton-Raphon's method.

Specifically, let $g(\delta) = \sum_{i=1}^{n}\left[\frac{nV_{xi}^2}{\left(\sum_{i=1}^{n}\frac{V_{xi}^2}{\lambda_{xi}+\delta}\right)(\lambda_{xi}+\delta)^2} - \frac{1}{\lambda_{xi}+\delta}\right]$, then with the current $\delta_n$, we update it by

$\delta_{n+1} = \delta_n - \frac{g(\delta_n)}{g'(\delta_n)}$ until $|\delta_{n+1} - \delta_n| \leq 10^{-6}$, say. Then the converged $\delta_{n+1}$ will be the MLE of $\delta$.

Finally, plugging the MLEs $\hat{\sigma}_g^2$ and $\hat{\sigma}_e^2$ back into (10) gives the estimated decorrelation matrix

$$\widehat{D}_x = \left(\hat{\sigma}_g^2 \Lambda_x + \hat{\sigma}_e^2 I\right)^{-\frac{1}{2}} U_x^T. \tag{26}$$

#### S3 Appendix: Proof of the soundness of the decorrelation procedure

Since $K_x$ is assumed of full rank, by eigen-decomposition we denote $K_x = U_x \Lambda_x U_x^T$, where $\Lambda_x = diag(\lambda_{x1}, \dots, \lambda_{xn}) \in \mathbb{R}^{n\times n}$ is a diagonal matrix with diagonal elements the eigenvalues $\lambda_{xi}$'s of $K_x$ in decreasing order, $U_x \in \mathbb{R}^{n\times n}$ is the eigenvector matrix with each column the eigenvector associated with the corresponding eigenvalues such that $U_x U_x^T = U_x^T U_x = I$. We assume $(\sigma_g^2 \Lambda_x + \sigma_e^2 I)$ is of full rank. Then for all the three analytical models, (1) for GWAS, (4) for emGWAS and (6) for TWAS, we have the following claim on their decorrelation matrix $D_x$.

**Claim 2.** *If*

$$D_x = \left(\sigma_g^2 \Lambda_x + \sigma_e^2 I\right)^{-\frac{1}{2}} U_x^T, \tag{10}$$

*then* $D_x(u+\varepsilon) \sim N(0, I)$.

*Proof.* For the LMMs given in (1), (4), and (6), the covariance matrix of the $n$ dimensional phenotype vector Y is

$$\begin{aligned} Var(Y) &= \sigma_g^2 K_x + \sigma_e^2 I \\ &= \sigma_g^2 U_x \Lambda_x U_x^T + \sigma_e^2 U_x U_x^T \\ &= U_x\left(\sigma_g^2 \Lambda_x + \sigma_e^2 I\right) U_x^T. \end{aligned}$$

With $D_x = \left(\sigma_g^2 \Lambda_x + \sigma_e^2 I\right)^{-\frac{1}{2}} U_x^T$, we have

$$\begin{aligned} Var(D_x Y) &= D_x Var(Y) D_x^T \\ &= D_x U\left(\sigma_g^2 \Lambda_x + \sigma_e^2 I\right) U_x^T D_x^T \\ &= \left(\sigma_g^2 \Lambda_x + \sigma_e^2 I\right)^{-\frac{1}{2}} U_x^T U_x\left(\sigma_g^2 \Lambda_x + \sigma_e^2 I\right) U_x^T \left(\left(\sigma_g^2 \Lambda_x + \sigma_e^2 I\right)^{-\frac{1}{2}} U_x^T\right)^T \\ &= \left(\sigma_g^2 \Lambda_x + \sigma_e^2 I\right)^{-\frac{1}{2}} U_x^T U_x\left(\sigma_g^2 \Lambda_x + \sigma_e^2 I\right) U_x^T U_x\left(\sigma_g^2 \Lambda_x + \sigma_e^2 I\right)^{-\frac{1}{2}} \\ &= \left(\sigma_g^2 \Lambda_x + \sigma_e^2 I\right)^{-\frac{1}{2}} I\left(\sigma_g^2 \Lambda_x + \sigma_e^2 I\right) I\left(\sigma_g^2 \Lambda_x + \sigma_e^2 I\right)^{-\frac{1}{2}} \\ &= \left(\sigma_g^2 \Lambda_x + \sigma_e^2 I\right)^0 \\ &= I. \end{aligned}$$

Hence the result.

#### S4 Appendix: Proof of the NCP in an LMM

For LMM (1), in order to identify the significant genetic variants, if we left multiply each term in (1) by the decorrelation matrix $D_x$ given by (10), then for the $j^{th}$ genetic variant vector, model (1) is reduced to

$$Y^* = \beta_{j0} X_0 + \beta_{j1} X_j^* + \varepsilon^*, \tag{27}$$

Where $Y^* = D_x Y = (Y_1^*, Y_2^*, \dots, Y_n^*)^T$ is the transformed phenotype vector, $X_0 = D_x \mathbf{1} = (D_{x1\cdot}, D_{x2\cdot}, \dots, D_{x\mathrm{n}\cdot})^T$ with $D_{x\mathrm{i}\cdot} = \sum_{j=1}^n D_{xij}$, $X_j^* = D_x X_j = (X_{1j}^*, X_{2j}^*, \dots, X_{nj}^*)^T$ is the transformed $j^{th}$ genetic variant vector, and $\varepsilon^* = D_x(u + \varepsilon)$ is the random vector such that $\varepsilon^* \sim N(0, I)$.

**Claim 3.** *For GWAS (1), the MLEs of $\beta_{j0}$ and $\beta_{j1}$ are*

$$\beta_{j1\_mle} = \frac{\sum_{i=1}^n X_{ij}^* Y_i^* \sum_{i=1}^n \mathrm{D}_{xi\cdot}^2 - \sum_{i=1}^n Y_i^* D_{xi\cdot} \sum_{i=1}^n X_{ij}^* D_{xi\cdot}}{\sum_{i=1}^n \left(X_{ij}^*\right)^2 \sum_{i=1}^n \mathrm{D}_{xi\cdot}^2 - \left(\sum_{i=1}^n X_{ij}^* \mathrm{D}_{xi\cdot}\right)^2}, \tag{28}$$

$$\beta_{j0\_mle} = \frac{\sum_{i=1}^n Y_i^* \mathrm{D}_{xi\cdot}}{\sum_{i=1}^n \mathrm{D}_{xi\cdot}^2} - \frac{\beta_{j1_{-mle}} \sum_{i=1}^n X_{ij}^* \mathrm{D}_{xi\cdot}}{\sum_{i=1}^n \mathrm{D}_{xi\cdot}^2}, \tag{29}$$

*And the variance of $\beta_{j1\_mle}$ is*

$$Var(\beta_{j1_{-mle}}) = \frac{\sum_{i=1}^n \mathrm{D}_{xi\cdot}^2}{\sum_{i=1}^n \left(X_{ij}^*\right)^2 \sum_{i=1}^n \widehat{D}_{xi\cdot}^2 - \left(\sum_{i=1}^n X_{ij}^* D_{xi\cdot}\right)^2}. \tag{30}$$

*Proof.* For LMM (1), or equivalently the transformed model (27), the log-likelihood function is

$$l(\beta_{j0};\beta_{j1}) = \ln\left(\prod_{i=1}^{n} f_{Y^*}\,(Y_1^*, Y_2^*, \cdots, Y_n^*)\right)$$

$$= -n\ln\sqrt{2\pi} - \sum_{i=1}^{n}\frac{\left(Y_i^* - \beta_{j0}\mathrm{D}_{xi\cdot} - \beta_{j1}X_{ij}^*\right)^2}{2}.$$

Then the MLE estimating equations for $\beta_{j0}$ and $\beta_{j1}$ are:

$$\frac{\partial}{\partial\beta_{j0}} l\left(\beta_{j0}, \beta_{j1}\right) = \sum_{i=1}^{n} \mathrm{D}_{xi\cdot}\left(Y_i^* - \beta_{j0}\mathrm{D}_{xi\cdot} - \beta_{j1}X_{ij}^*\right) = \sum_{i=1}^{n} Y_i^*\mathrm{D}_{xi\cdot} - \beta_{j0}\sum_{i=1}^{n}\mathrm{D}_{xi\cdot}^2 - \beta_{j1}\sum_{i=1}^{n} X_{ij}^*\mathrm{D}_{xi\cdot} = 0, (31)$$

$$\frac{\partial}{\partial\beta_{j1}} l\left(\beta_{j0}, \beta_{j1}\right) = \sum_{i=1}^{n} X_{ij}^*\left(Y_i^* - \mathrm{D}_{xi\cdot}\beta_{j0} - \beta_{j1}X_{ij}^*\right) = \sum_{i=1}^{n} X_{ij}^*\,Y_i^* - \beta_{j0}\sum_{i=1}^{n} X_{ij}^*\,\mathrm{D}_{xi\cdot} - \beta_{j1}\sum_{i=1}^{n}\left(X_{ij}^*\right)^2 = 0. (32)$$

From (31) we obtain (29). Plugging (29) into (32) gives:

$$0 = \sum_{i=1}^{n} X_{ij}^*\,Y_i^* - \frac{\sum_{i=1}^{n} Y_i^*\mathrm{D}_{xi\cdot}\,\sum_{i=1}^{n} X_{ij}^*\,\mathrm{D}_{xi\cdot}}{\sum_{i=1}^{n}\mathrm{D}_{xi\cdot}^2} + \frac{\beta_{j1}(\sum_{i=1}^{n} X_{ij}^*\mathrm{D}_{xi\cdot})^2}{\sum_{i=1}^{n}\mathrm{D}_{xi\cdot}^2} - \beta_{j1}\sum_{i=1}^{n}\left(X_{ij}^*\right)^2$$

$$= \frac{\sum_{i=1}^{n} X_{ij}^*\,Y_i^*\sum_{i=1}^{n}\mathrm{D}_{xi\cdot}^2 - \sum_{i=1}^{n} Y_i^*\,\mathrm{D}_{xi\cdot}\,\sum_{i=1}^{n} X_{ij}^*\,\mathrm{D}_{xi\cdot}}{\sum_{i=1}^{n}\mathrm{D}_{xi\cdot}^2} - \beta_{j1}\frac{\sum_{i=1}^{n}\left(X_{ij}^*\right)^2\sum_{i=1}^{n}\mathrm{D}_{xi\cdot}^2 - \left(\sum_{i=1}^{n}\mathrm{D}_{xi\cdot}\,X_{ij}^*\right)^2}{\sum_{i=1}^{n}\mathrm{D}_{xi\cdot}^2},$$

and thus

$$\beta_{j1_{\mathrm{mle}}} = \frac{\sum_{i=1}^{n} X_{ij}^*\,Y_i^*\sum_{i=1}^{n}\mathrm{D}_{xi\cdot}^2 - \sum_{i=1}^{n} Y_i^*\,D_{xi\cdot}\,\sum_{i=1}^{n} X_{ij}^*\,D_{xi\cdot}}{\sum_{i=1}^{n}\left(X_{ij}^*\right)^2\sum_{i=1}^{n}\mathrm{D}_{xi\cdot}^2 - \left(\sum_{i=1}^{n} X_{ij}^*\mathrm{D}_{xi\cdot}\right)^2}$$

$$= \frac{\sum_{i=1}^{n} \mathrm{D}_{xi\cdot}^2 \sum_{i=1}^{n} \left( X_{ij}^* - D_{xi\cdot} \frac{\sum_{i=1}^{n} X_{ij}^* \mathrm{D}_{xi\cdot}}{\sum_{i=1}^{n} \mathrm{D}_{xi\cdot}^2} \right) Y_i^*}{\sum_{i=1}^{n} \left(X_{ij}^*\right)^2 \sum_{i=1}^{n} \mathrm{D}_{xi\cdot}^2 - \left( \sum_{i=1}^{n} \mathrm{D}_{xi\cdot} X_{ij}^* \right)^2},$$

i.e. (28) holds. Since by Claim 2 we have $Var(D_x Y) = Var(Y^*) = I$, direct calculation gives the variance of $\beta_{j1\_mle}$ as

$$Var\left(\beta_{j1\_mle}\right) = \left( \frac{\sum_{i=1}^{n} D_{xi\cdot}^2}{\sum_{i=1}^{n} \left(X_{ij}^*\right)^2 \sum_{i=1}^{n} D_{xi\cdot}^2 - \left( \sum_{i=1}^{n} X_{ij}^* D_{xi\cdot} \right)^2} \right)^2 Var\left( \sum_{i=1}^{n} \left( X_{ij}^* - D_{xi\cdot} \frac{\sum_{i=1}^{n} X_{ij}^* D_{xi\cdot}}{\sum_{i=1}^{n} \mathrm{D}_{xi\cdot}^2} \right) Y_i^* \right)$$

$$= \left( \frac{\sum_{i=1}^{n} \mathrm{D}_{xi\cdot}^2}{\sum_{i=1}^{n} \left(X_{ij}^*\right)^2 \sum_{i=1}^{n} \mathrm{D}_{xi\cdot}^2 - \left( \sum_{i=1}^{n} X_{ij}^* D_{xi\cdot} \right)^2} \right)^2 \sum_{i=1}^{n} \left( X_{ij}^* - \mathrm{D}_{xi\cdot} \frac{\sum_{i=1}^{n} X_{ij}^* D_{xi\cdot}}{\sum_{i=1}^{n} \mathrm{D}_{xi\cdot}^2} \right)^2 Var(Y_i^*)$$

$$= \frac{\sum_{i=1}^{n} \left( X_{ij}^* \sum_{i=1}^{n} \mathrm{D}_{xi\cdot}^2 - \mathrm{D}_{xi\cdot} \sum_{i=1}^{n} X_{ij}^* \mathrm{D}_{xi\cdot} \right)^2}{\left( \sum_{i=1}^{n} \left(X_{ij}^*\right)^2 \sum_{i=1}^{n} \mathrm{D}_{xi\cdot}^2 - \left( \sum_{i=1}^{n} X_{ij}^* D_{xi\cdot} \right)^2 \right)^2}$$

$$= \frac{\sum_{i=1}^{n} \left[ \left(X_{ij}^*\right)^2 \left( \sum_{i=1}^{n} \mathrm{D}_{xi\cdot}^2 \right)^2 - 2 X_{ij}^* \mathrm{D}_{xi\cdot} \sum_{i=1}^{n} \mathrm{D}_{xi\cdot}^2 \sum_{i=1}^{n} X_{ij}^* \mathrm{D}_{xi\cdot} + \mathrm{D}_{xi\cdot}^2 \left( \sum_{i=1}^{n} X_{ij}^* \mathrm{D}_{xi\cdot} \right)^2 \right]}{\left( \sum_{i=1}^{n} \left(X_{ij}^*\right)^2 \sum_{i=1}^{n} \mathrm{D}_{xi\cdot}^2 - \left( \sum_{i=1}^{n} X_{ij}^* D_{xi\cdot} \right)^2 \right)^2}$$

$$= \frac{\sum_{i=1}^{n} \left(X_{ij}^*\right)^2 \left( \sum_{i=1}^{n} \mathrm{D}_{xi\cdot}^2 \right)^2 - 2 \sum_{i=1}^{n} \widehat{D}_{xi\cdot}^2 \left( \sum_{i=1}^{n} X_{ij}^* \mathrm{D}_{xi\cdot} \right)^2 + \sum_{i=1}^{n} \mathrm{D}_{xi\cdot}^2 \left( \sum_{i=1}^{n} X_{ij}^* D_{xi\cdot} \right)^2}{\left( \sum_{i=1}^{n} \left(X_{ij}^*\right)^2 \sum_{i=1}^{n} \widehat{D}_{xi\cdot}^2 - \left( \sum_{i=1}^{n} X_{ij}^* D_{xi\cdot} \right)^2 \right)}$$

$$= \frac{\sum_{i=1}^{n} \left(X_{ij}^*\right)^2 \left( \sum_{i=1}^{n} \mathrm{D}_{xi\cdot}^2 \right)^2 - \sum_{i=1}^{n} \mathrm{D}_{xi\cdot}^2 \left( \sum_{i=1}^{n} X_{ij}^* D_{xi\cdot} \right)^2}{\left( \sum_{i=1}^{n} \left(X_{ij}^*\right)^2 \sum_{i=1}^{n} \mathrm{D}_{xi\cdot}^2 - \left( \sum_{i=1}^{n} X_{ij}^* D_{xi\cdot} \right)^2 \right)^2}$$

$$= \frac{\sum_{i=1}^{n} \mathrm{D}_{xi\cdot}^2}{\sum_{i=1}^{n} \left(X_{ij}^*\right)^2 \sum_{i=1}^{n} \widehat{D}_{xi\cdot}^2 - \left( \sum_{i=1}^{n} X_{ij}^* D_{xi\cdot} \right)^2}.$$

Since the true variance components $\sigma_g^2$ and $\sigma_e^2$ are unknown and thus the decorrelation matrix $D_x$ is unknown either, we use the estimated $\widehat{D}_x$ in (26) instead. Plugging $\widehat{D}_x$ into (28) and (30), we obtain an estimated MLE and an estimated standard deviation of the MLE given respectively by

$$\hat{\beta}_{j1\_mle} = \frac{\sum_{i=1}^{n} \hat{X}_{ij}^{*} \hat{Y}_{i}^{*} \sum_{i=1}^{n} \widehat{\mathrm{D}}_{xi\cdot}^{2} - \sum_{i=1}^{n} \hat{Y}_{i}^{*} \widehat{D}_{xi\cdot} \sum_{i=1}^{n} \hat{X}_{ij}^{*} \widehat{D}_{xi\cdot}}{\sum_{i=1}^{n} \left(\hat{X}_{ij}^{*}\right)^{2} \sum_{i=1}^{n} \widehat{\mathrm{D}}_{xi\cdot}^{2} - \left(\sum_{i=1}^{n} \hat{X}_{ij}^{*} \widehat{\mathrm{D}}_{xi\cdot}\right)^{2}},$$

$$\widehat{SD}\left(\beta_{j1\_mle}\right) = \sqrt{\frac{\sum_{i=1}^{n} \widehat{D}_{xi\cdot}^{2}}{\sum_{i=1}^{n} \left(X_{ij}^{*}\right)^{2} \sum_{i=1}^{n} \widehat{D}_{xi\cdot}^{2} - \left(\sum_{i=1}^{n} \widehat{D}_{xi\cdot} X_{ij}^{*}\right)^{2}}},$$

where $\hat{X}_{j}^{*} = \widehat{D}_{x} X_{j} = (\hat{X}_{1j}^{*}, \hat{X}_{2j}^{*}, \dots, \hat{X}_{nj}^{*})^{T}$, $\hat{Y}^{*} = \widehat{D}_{x} Y = (\hat{Y}_{1}^{*}, \hat{Y}_{2}^{*}, \dots, \hat{Y}_{n}^{*})^{T}$, and $\widehat{D}_{x\mathrm{i}\cdot} = \sum_{j=1}^{n} \widehat{D}_{xij}$.

Recall the NCP pf the $t$ test statistic for the slope $\beta$ in a simple linear regression model is given by $\upsilon = \frac{\beta}{SD(\hat{\beta})}$. Then the NCP $\upsilon_{Gj}$ based on MLE under the true alternative $\beta_{j1}$ is given by

$$\upsilon_{Gj} = \frac{\beta_{j1}}{SD(\beta_{j1\_mle})} = \frac{\beta_{j1} \sqrt{\sum_{i=1}^{n} \left(X_{ij}^{*}\right)^{2} \sum_{i=1}^{n} \mathrm{D}_{xi\cdot}^{2} - \left(\sum_{i=1}^{n} X_{ij}^{*} D_{xi\cdot}\right)^{2}}}{\sqrt{\sum_{i=1}^{n} \mathrm{D}_{xi\cdot}^{2}}}. \tag{33}$$

Since $\beta_{j1}$ and $D_{x}$ are unknown, when replacing them with the estimated MLE $\hat{\beta}_{j1\_mle}$ and $\widehat{D}_{x}$ in (26) respectively into (33), an estimated NCP $\hat{\upsilon}_{Gj}$ for GWAS is given by (11).

#### S5 Appendix: Simulating expression and phenotype that have prespecified variance components under the causality scenario

For a certain $l^{th}$ selected gene $\left(l = 1,2, \cdots, n_{z-sig}\right)$, the gene expression $Z_{(l)}$ was simulated by its corresponding local $n_{z(l)-sig}$ randomly picked genetic variants.

$$Z_{(l)} = \left( \frac{1}{Var\left(\sum_{k=1}^{n_{z(l)-sig}} X_{1(l)(k)} b_{(l)(k)}\right) + Var(\varepsilon_{(l)})} \right)^{\frac{1}{2}} \left(X_{(l)} b_{(l)} + \varepsilon_{(l)}\right), \tag{34}$$

where the $n \times 1$ vector $Z_{(l)}$ is the $l^{th}$ significant gene expression of $n$ individuals, $X_{(l)}$ is an $n \times n_{z(l)-sig}$ matrix of randomly picked $n_{z(l)-sig}$ qualified variants in the $l^{th}$ significant gene, $X_{i(l)(k)}$ denotes the $k^{th}$ causal variant in the $l^{th}$ causal gene for the $i^{th}$ individual with $i = 1, \ldots, n$ , $b_{(l)} = (b_{(l)(1)}, b_{(l)(2)}, \ldots, b_{(l)(n_{z(l)-sig})})^T$ is a $n_{z(l)-sig} \times 1$ vector of effect sizes of the causal variants, $\varepsilon_{(l)}$ is an $n \times 1$ vector of errors such that $\varepsilon_{(l)} \sim N(0,\mathrm{Var}(\varepsilon_{(l)})$. In our simulation $b_{(l)}$ was generated with each element a random value between 0 and 1. The gene expression was simulated based on the contribution from $X_{(l)} b_{(l)}$ and residuals from the expression heritability $h^2_{\mathrm{x=>z}}$. The expression heritability quantifies the contribution of genetic variants to the gene expression by

$$h^2_{\mathrm{x=>z}} = \frac{Var\left(\sum_{k=1}^{n_{z(l)-sig}} X_{1(l)(k)} b_{(l)(k)}\right)}{Var\left(\sum_{k=1}^{n_{z(l)-sig}} X_{1(l)(k)} b_{(l)(k)}\right) + Var(\varepsilon_{(l)})}, \tag{35}$$

where $Var\left(\sum_{k=1}^{n_{z(l)-sig}} X_{1(l)(k)} b_{(l)(k)}\right)$ was estimated by the sample variance given by

$$\widehat{Var}\left(\sum_{k=1}^{n_{z(l)-sig}} X_{1(l)(k)} b_{(l)(k)}\right) = \frac{1}{n-1} \sum_{i=1}^{n} \left( \sum_{k=1}^{n_{z(l)-sig}} X_{i(l)(k)} b_{(l)(k)} - \frac{1}{n} \sum_{i=1}^{n} \sum_{k=1}^{n_{z(l)-sig}} X_{i(l)(k)} b_{(l)(k)} \right)^2. \tag{36}$$

Plugging the estimated variance (36) into (35), and estimate of $Var(\varepsilon_{(l)})$ was given by

$$\widehat{Var}(\varepsilon_{(l)}) = \left( \frac{1}{h^2_{x=>\mathrm{z}}} - 1 \right) \widehat{Var}\left( \sum_{k=1}^{n_{z(l)-sig}} X_{1(l)(k)} b_{(l)(k)} \right). \tag{37}$$

Then the errors $\varepsilon_{(l)}$ are generated randomly from the normal distribution $N\sim\left(0,\widehat{Var}\left(\varepsilon_{(l)}\right)I\right)$. After replacing $Var\left(\sum_{k=1}^{n_{z(l)-sig}} X_{1(l)(k)}b_{(l)(k)}\right)$ and $Var(\,\varepsilon_{(l)})$ in model (34) with their estimates given in (36) and (37) respectively, we can then generate $Z_{(l)}$.

Following the steps above, the $n_{z-sig}$ causal genes' expression $Z_{(l)}$'s are simulated. Since in causality scenario gene expression explains phenotypes, the latter were generated from the $n_{z-sig}$ causal genes. More specifically,

$$Y_{ca} = \left(\frac{1}{Var\left(\sum_{l=1}^{n_{z-sig}} Z_{1(l)}b_{c(l)}\right) + Var(\,\varepsilon_c)}\right)^{\frac{1}{2}} (Zb_c + \varepsilon_c), \quad (38)$$

where the $n \times 1$ vector $Y_{ca}$ is the phenotypes under causality scenario, $Z = (Z_{(1)}, Z_{(2)}, \dots, Z_{(n_{z-sig})})$ is an $n \times n_{z-sig}$ matrix of significant genes' expressions, $b_c = (b_{c(1),} b_{c(2),} \dots, b_{c(n_{z-sig})})^T$ is an $n_{z-sig} \times 1$ vector of effect sizes of the gene expressions, and the $n \times 1$ vector of errors $\varepsilon_c \sim N(0, Var(\varepsilon_c))$. Similarly, $b_c$ was generated with each element a random value between 0 and 1. We estimated $Var\left(\sum_{l=1}^{n_{z-sig}} Z_{(l)}b_{c(l)}\right)$ by sample variance

$$\widehat{Var}\left(\sum_{l=1}^{n_{z-sig}} Z_{1(l)}b_{c(l)}\right) = \frac{1}{n-1}\sum_{i=1}^{n}\left(\sum_{l=1}^{n_{z-sig}} Z_{i(l)}b_{c(l)} - \frac{1}{n}\sum_{i=1}^{n}\sum_{l=1}^{n_{z-sig}} Z_{i(l)}b_{c(l)}\right)^2. \quad (39)$$

Following the relationship between PVX and the variance component, i.e.

$$h^2_{z=>y} = \frac{Var\left(\sum_{l=1}^{n_{z-sig}} Z_{1(l)} b_{c(l)}\right)}{Var\left(\sum_{l=1}^{n_{z-sig}} Z_{1(l)} b_{c(l)}\right) + Var(\varepsilon_c)} \quad (40)$$

$Var(\varepsilon_c)$ can be thus estimated by

$$\widehat{Var}(\varepsilon_c) = \left(\frac{1}{h^2_{z=>y}} - 1\right) \widehat{Var}\left(\sum_{l=1}^{n_{z-sig}} Z_{1(l)} b_{c(l)}\right). \quad (41)$$

Then the errors $\varepsilon_{ca}$ were generated randomly from the normal distribution $N \sim \left(0, \widehat{Var}(\varepsilon_c)\right)$. Finally we replaced $Var\left(\sum_{l=1}^{n_{z-sig}} Z_{1(l)} b_{c(l)}\right)$ and $Var(\varepsilon_c)$ in model (38) with their estimates given in (39) and (41) respectively, and then $Y_{ca}$ in (38) can be simulated.

**S6 Appendix: Simulating expression and phenotype that have prespecified variance components under the pleiotropy scenario**

Similar to the process in **S5 Appendix**, the phenotypes were simulated from a combination of $n_{x-sig}$ significant genetic variants directly as follow

$$Y_p = \left(\frac{1}{Var\left(\sum_{k=1}^{n_{x-sig}} X_{1(k)} b_{p(k)}\right) + Var(\varepsilon_p)}\right)^{1/2} \left(\mathrm{X} b_p + \varepsilon_p\right), \quad (42)$$

where the $n \times 1$ vector $Y_p$ is the phenotypes under causality scenario, $X = (X_{(1)}, X_{(2)}, \ldots, X_{(n_{x-sig})})$ is an $n \times n_{x-sig}$ matrix of significant genetic variants, $b_p$ is a $n_{x-sig} \times 1$ vector of effect sizes, and $\varepsilon_p$ is an $n \times 1$ vector of residuals. Similarly, we first generated the slopes $b_p$ randomly with each element between 0 and 1, then we obtained $\mathrm{X} b_p$ and further the sample variance

$$\widehat{Var}\left(\sum_{k=1}^{n_{x-sig}} X_{1(k)} b_{p(k)}\right) = \frac{1}{n-1}\sum_{i=1}^{n}\left(\sum_{k=1}^{n_{x-sig}} X_{i(k)} b_{p(k)} - \frac{1}{n}\sum_{i=1}^{n}\sum_{k=1}^{n_{x-sig}} X_{i(k)} b_{p(k)}\right)^2. \quad (43)$$

Based on the prespecified trait heritability $h^2_{\mathrm{x=>y}}$ and its relationship with variance components given by

$$h^2_{\mathrm{x=>y}} = \frac{Var\left(\sum_{k=1}^{n_{x-sig}} X_{1(k)} b_{p(k)}\right)}{Var\left(\sum_{k=1}^{n_{x-sig}} X_{1(k)} b_{p(k)}\right) + Var(\varepsilon_p)}, \quad (44)$$

$Var(\varepsilon_p)$ can then be estimated by

$$\widehat{Var}(\varepsilon_p) = \left(\frac{1}{h^2_{x=>\mathrm{y}}} - 1\right)\widehat{Var}\left(\sum_{k=1}^{n_{x-sig}} X_{1(k)} b_{p(k)}\right). \quad (45)$$

Then the errors $\varepsilon_p$ were generated randomly from the normal distribution $N\sim\left(0, \widehat{Var}(\varepsilon_p)\right)$. Finally we replaced $Var\left(\sum_{k=1}^{n_{x-sig}} X_{1(k)} b_{p(k)}\right)$ and $Var(\varepsilon_p)$ in model (42) with their estimates given in (43) and (45) respectively, and then $Y_p$ in (42) can be simulated.
